# Auxin-responsive (phospho)proteome analysis reveals regulation of cell cycle and ethylene signaling during rice crown root development

**DOI:** 10.1101/2021.03.30.437660

**Authors:** Harshita Singh, Zeenu Singh, Tingting Zhu, Xiangyu Xu, Bhairavnath Waghmode, Tushar Garg, Shivani Yadav, Debabrata Sircar, Ive De Smet, Shri Ram Yadav

## Abstract

The rice root system, which primarily consists of adventitious/crown roots (AR/CR) developed from the coleoptile base, is an excellent model system for studying shoot-to-root trans-differentiation process. We reveal global changes in protein and metabolite abundance, and protein phosphorylation in response to an auxin stimulus during CR development. Global proteome and metabolome analyses of developing crown root primordia (CRP) and emerged CRs uncovered that the biological processes associated with chromatin conformational change, gene expression, and cell cycle were translationally regulated by auxin signaling. Spatial gene expression pattern analysis of differentially abundant proteins disclosed their stage-specific dynamic expression pattern during CRP development. Further, our tempo-spatial gene expression and functional analyses revealed that auxin creates a regulatory feedback module during CRP development and activates ethylene biosynthesis exclusively during CRP initiation. Ethylene signaling promotes CR formation by repressing the cytokinin response regulator, *OsRR2*. Additionally, the (phospho)proteome analysis identified differential phosphorylation of the Cyclin-dependent kinase G-2 (OsCDKG;2), and cell wall proteins, in response to auxin signaling, suggesting that auxin-dependent phosphorylation may be required for cell cycle activation, and cell wall synthesis during root organogenesis. Thus, our study provides evidence for the translational and post-translational regulation during CRP trans-differentiation downstream of the auxin signaling pathway.

**Highlight:** Global (phospho)proteome and metabolic profiling of rice CRP and CRs uncover differential proteins and metabolites associated with gene expression, cell cycle, ethylene signaling and cell wall synthesis during CR development.

## Introduction

The root architecture of plants critically determines their nutrient and water absorption efficiency. The root system of monocot cereal plants is composed of seminal embryonic axile roots from the radicle and axile adventitious roots from the coleoptile nodes (Mai *et al.*, 2014). In rice, adventitious roots arise constitutively from the basal coleoptile/stem nodes in a circular pattern, also known as nodal or crown roots (CR). These post-embryonic shoot borne CRs dominate the mature root system of the rice plant. During rice CR formation, positional cues establish the developmental program for crown root primordium (CRP) initiation inside the coleoptile base, which leads to an emerged CR from the nodes. The CR founder cells originate from the ground meristem tissues that are positioned peripheral to the vascular tissues (Itoh *et al.*, 2005). The founder cells re-activate cell division to form three layers of fundamental tissues in a dome-shaped CRP. During later stages of CRP outgrowth, cell differentiation and tissue patterning take place. Finally, vacuolation, cell elongation, and cell wall remodelling lead to the rupture of the epidermal layer to facilitate emergence of CRP (Itoh *et al.*, 2005). Root development is a cumulative effect of action and interaction of different hormonal signaling pathways and transcription factors in plants (Chaiwanon *et al.*, 2016). Auxin is a key regulator of root organogenesis in plants and auxin maxima is a prerequisite for CRP initiation. Besides CRP initiation, it also regulates later stages of CR development, such as CRP outgrowth and CR emergence. A gradient of auxin is established in specific ground meristem cells, via local auxin biosynthesis and polar auxin transport, conjugation, and degradation (Chapman and Estelle, 2009). Auxin biosynthetic genes of the *OsYUCCA* gene family, and regulators of polar auxin transport (*CROWN-ROOTLESS4/OsGNOM1; OsCRL4/OsGNOM1* and *PINFORMED1; OsPIN),* positively regulate CR number in rice (Xu *et al.*, 2005; Yamamoto *et al.*, 2007). Additionally, the gain-of-function mutant *Osiaa23*, a negative regulator of auxin signaling, shows impaired QC maintenance and defects in CR initiation (Jun *et al.*, 2011). Furthermore, the *CULLIN-ASSOCIATED AND NEDDYLATION- DISSOCIATED1 (OsCAND1)*, a gene required for SCF^TIR1^ activity during activation of auxin signaling, regulates G2/M cell cycle transition plays a crucial role in CR emergence (Wang *et al.*, 2011). Additionally, transcription factors and other genes such as *ADVENTITIOUS ROOTLESS 1 (ARL1)/CROWN ROOTLESS 1 (CRL1), CROWN ROOTLESS 5 (CRL5), AP2/ETHYLENE-RESPONSIVE FACTOR3 (ERF3), NARROW LEAF 1 (NAL1), CROWN ROOTLESS 4 (CRL4)/OsGNOM1, CHROMATIN REMODELING 4 (CHR4)/CROWN ROOTLESS 6 (CRL6), SQUAMOSA PROMOTER BINDING PROTEIN-LIKE3 (OsSPL3), WUSCHEL-related HOMEOBOX 10 (OsWOX10), OsWOX11*, and *AP2/ETHYLENE RESPONSE FACTOR 40 (OsAP2/ERF-40)* also regulate CR development (Inukai *et al.*, 2005; Kitomi *et al.*, 2008, 2011; Zhao *et al.*, 2009; 2015; Wang *et al.*, 2016; Neogy *et al.*, 2019; Shao *et al.*, 2019; Garg et al., 2020).

Auxin signaling is activated by post-translational degradation of Aux/IAA repressor proteins that otherwise inhibit the activity of auxin response factors (ARFs) (Chapman and Estelle, 2009). Recently, transcript profiling was used to decipher the transcriptional gene regulatory network controlling CR development downstream of auxin signaling pathway (Neogy *et al.*, 2019). However, there is no linear correlation between transcript and protein abundance (Vogel and Marcotte, 2012; Liu *et al.*, 2016). Therefore, it becomes relevant to use proteomics-based approaches to determine the global proteome to reveal translational regulations. Importantly, cellular signaling involves reversible post-translational protein modifications as a switch to regulate protein activity and localization. Phosphorylation and dephosphorylation contribute to a robust mechanism for such a switch in eukaryotic cells and play a vital role in regulating signal transduction pathways by modulating protein-protein interactions and protein activity (Mithoe and Menke, 2011). In plants, phosphorylation-mediated signaling is the center of many developmental and physiological processes, hormonal responses and stress-related signaling. Understanding the detailed contribution of proteins and their post-translational modifications during cellular signaling pathways requires precise quantification methods. Mass spectrometry-based protein analysis is an essential tool for the identification of numerous peptides and phosphopeptides in plant tissues (Li *et al.*, 2015*a*). The present study investigated auxin-mediated modulation in the proteome and metabolome during rice CRP development and the growth of emerged CRs. Using a label-free, quantitative (phospho)proteomics-based approach (Vu *et al.*, 2016), we have identified proteins whose abundance is altered upon auxin treatment. Further, LCM-seq and RNA *in situ* hybridization together with use of pharmacological inhibitors of hormonal signaling pathways showed the biological processes such as cell cycle and auxin and ethylene signaling pathways are translationally regulated during CRP trans-differentiation. Moreover, we also revealed phosphorylation-based protein modification of cell cycle and cell wall associated proteins in response to auxin signaling during CR development.

## Materials and methods

### Plant material, growth conditions and treatments

The rice (*Oryza sativa indica*) IR64 seeds were surface sterilised and germinated in the half-strength liquid Hoagland media (Himedia Pvt Ltd, India) under hydroponics conditions at 28°C with the 16 h photoperiod. To deplete the endogenous auxin, six-day old seedlings with uniform growth were removed from the hydroponics and transferred to freshly prepared KPSC buffer containing 2% sucrose, 10mM potassium phosphate (pH 6) and 50 μM chloramphenicol for 8 hours. The buffer was replaced after every one hour (Thakur *et al.*, 2001). Later the seedlings transferred to fresh KPSC buffer supplemented with 10 μM IAA and DMSO as control for 3h for (phospho)proteome and 12h for metabolome analyses. About 2 mm coleoptile bases and all crown roots of the seedlings were collected and frozen in liquid nitrogen.

### Protein extraction, tryptic digestion, phosphopeptide enrichment and LC-MS/MS analysis

Total protein extraction was conducted on four biological replicate samples according to the described procedure with minor modifications (Vu *et al.*, 2016). The enrichment for the phosphopeptide procedure was performed as reported previously (Vu *et al.*, 2016). Ultimate 3000 RSLC nano LC was used to analyse samples via LC-MS/MS on an (Thermo Fisher Scientific, Bremen, Germany) in-line connected to a Q Exactive mass spectrometer (Thermo Fisher Scientific). Later, MS/MS spectra were searched against the rice database, downloaded from UniProt, and MS1-based label-free quantification was acquired with the MaxQuant software (version 1.5.4.1) from Orbitrap instruments (Cox and Mann, 2008; Cox *et al.*, 2014). The detailed procedure has been discussed in the Supplementary Methodology.

### Gene ontology analysis

Gene ontology enrichment analysis was performed using PLAZA 4.5 monocot workbench (Van Bel *et al.*, 2018). For the rice proteome datasets, 201 proteins with significant upregulation, 133 proteins with significant down-regulation and 3571 total identified proteomes were analyzed, using the dataset of the whole theoretical rice genome as a background model. Similarly, for the rice phosphoproteome dataset, 66 peptides with significant changes in phosphorylation abundance were analyzed, using the dataset of the whole theoretical rice genome as a background model. P-value cutoff was set at < 0.01 and only terms enriched in either condition were presented.

### Protein-protein interaction using STRINGv11

We analyzed protein-protein interaction among the differentially regulated protein dataset using STRINGv11(Szklarczyk *et al.*, 2019). To highlight highly confident interactions, the required confidence score was adjusted to > 0.90 for upregulated and downregulated protein network interactions. STRING protein-protein interaction prediction is based on the previous reports and data available for genomic homology, gene fusion, occurrence in the same metabolic pathways, co-expression, experiments, database and text mining. The scores of all the methods used for the protein-protein interaction prediction was used to calculate a combined score.

### Spatial expression pattern analysis

The expression pattern of genes during crown root initiation and crown root outgrowth was derived from the genome-wide high resolution expression dataset (Garg *et al.*, 2020). Genes with adjusted q value < 0.05 and log2 fold change ± 0.5 were considered differentially expressed. The expression pattern of genes in various organs and different root zones was obtained using CoNekT database (Proost and Mutwil, 2018) (https://conekt.sbs.ntu.edu.sg/heatmap/). The heatmaps were generated using Heatmapper tool (Babicki *et al.*, 2016) (http://heatmapper.ca/expression/).

### Metabolic profiling and data analysis

For metabolic profiling, rice seedlings were treated with IAA for 12h. About 100 mg fresh tissues were processed for GC-MS analysis as described earlier (Sarkate et al., 2021). Processing of GC–MS data, Multivariate Statistical Analysis, and Partial Least Squares Discriminant Analysis (PLS-DA) were performed as described by Kumar et al., 2021. Important features were extracted from PLS-DA data using Variable importance in projection (VIP) scores and heatmap was created using Metaboanalyst 4.0. A detailed method is described in supplementary methods.

### RNA in situ hybridization

To synthesise DIG-UTP-labeled riboprobes, the 174 bp gene-specific region of *OsRP1*, 170 bp gene-specific region of *OsACO1* and 183 bp gene-specific region of *OsFMO* were cloned into pBluescript SK+ as a blunt in EcoRV site. The sense probes for *OsRP1, OsACO1* and *OsFMO* were generated by linearising the pBluescript SK+ clones using HindIII and transcription with T3 RNA polymerase (Sigma-Aldrich). The anti-sense probes were synthesised using EcoRV linearised pBluescript SK+ clones, transcribed using T7 polymerase (Roche). Hybridization was performed on cross-sections as described by Neogy et al. (2020). Alkaline phosphatase-conjugated anti-DIG antibodies (Sigma-Aldrich) and NBT/BCIP substrate (Sigma-Aldrich) was used to develop signal and Entellan (Merck-Millipore, Darmstadt, Germany) for section mounting.

### Phenotyping

Rice Rice (*O. sativa* var IR-64) seeds were dehusked, surface sterilised and germinated on ½ MS media (Himedia) with 1% sucrose (Himedia) and 0.4% clerigel (Himedia) at 28°C with the 16h photoperiod. To study the effect of auxin signaling inhibitor on CR development, IR64 seeds were germinated on ½ MS media supplemented with 0.5 μM and 1 μM NPA (Sigma-Aldrich). The number of crown roots were measured on 6^th^ day post-germination. Similarly, to analyse the effect of ethylene inhibitors on CR number, IR64 seeds were germinated on media containing 50 μM AgNO_3_, 100 μM AgNO_3_. The number of crown roots were measured on 6^th^ day and 8^th^ day post-germination for AgNO_3_ experiment. Further, we treated the 3^rd^ day old seedling with 1ppm MCP, a competitive inhibitor of ethylene signaling and the CR number were calculated on 9^th^ day post-germination. The student t-test was performed on the calculated CR numbers using the data analysis tool in Microsoft excel.

### Sampling, RNA extraction, reverse transcription and real-time PCR

About 2 mm coleoptile base was collected from treated plants and RNA was extracted using Tri-Reagent (Sigma-Aldrich) following the manufacturer’s protocol. The RNA integrity was assured using agarose gel and quantity was measured using Nanodrop. The 10 μg of RNA was treated with DNaseI (New England BioLabs, USA), followed by phenol: chloroform treatment. To check the expression of key CR development regulators, 1μg of total RNA were used for cDNA synthesis using iScript™ cDNA Synthesis Kit (Bio-rad Laboratories, India). For quantitative real-time PCR (qRT-PCR), we used 10ng of cDNA, iTAQ Universal SYBR Green Supermix (Bio-rad, Laboratories, India) and 250 nM of gene-specific primers in QuantStudio 3 machine (Thermo Scientific). To calculate log_2_ fold change in treated and mock samples, ΔΔCt values were normalised by *UBQ5*. A list of primer sequences is provided as Supplementary Table S2.

## Results

### Global and auxin-regulated (phospho)proteome analysis during CR development

Previously, our RNA-seq based transcript profiling study revealed transcriptional regulation of auxin signaling during CRP development (Neogy *et al.*, 2019) but regulation at translational and post-translational protein modification levels was not yet studied. Thus, to reveal auxin-dependent regulation of protein levels and phosphorylation status during CR development, we treated wild-type seedlings with IAA and performed LC-MS/MS-based comparative (phospho)proteome analysis of the rice coleoptile base containing developing CRP and emerged CRs (Fig. 1A, B). The rationale behind using coleoptile bases and CRs was to cover all developmental stages of CR formation, from CRP initiation, to their outgrowth and the emerged CRs. Auxin induction in the IAA-treated samples was validated by analysing expression levels of a few known auxin-activated genes using quantitative real-time PCR (Fig. 1C). An overview of the comparative (phospho)proteome workflow is depicted in (Fig. 1D). The LC-MS/MS analysis of mock and auxin-treated rice coleoptile bases resulted in the identification of 3571 proteins (Supplementary Dataset S1). The values of average pearson correlation calculated using the data from replicates were 0.981 and 0.966 for mock and auxin treated samples, respectively, which demonstrates quantitative reproducibility of our proteome data (Supplementary Fig. S1). Statistical analysis (p < 0.05) determined that the abundance of 201 proteins was increased upon IAA treatment, whereas the abundance of 133 proteins was decreased in response to IAA treatment (Fig. 1E; Supplementary Dataset S1). Of these, 49 proteins were not detected in the mock-treated samples, but their protein levels were induced upon auxin treatment. In contrast, 19 proteins were present in the mock-treated samples, but they were not detected upon IAA treatment (Supplementary Dataset S1).

**Fig. 1.**
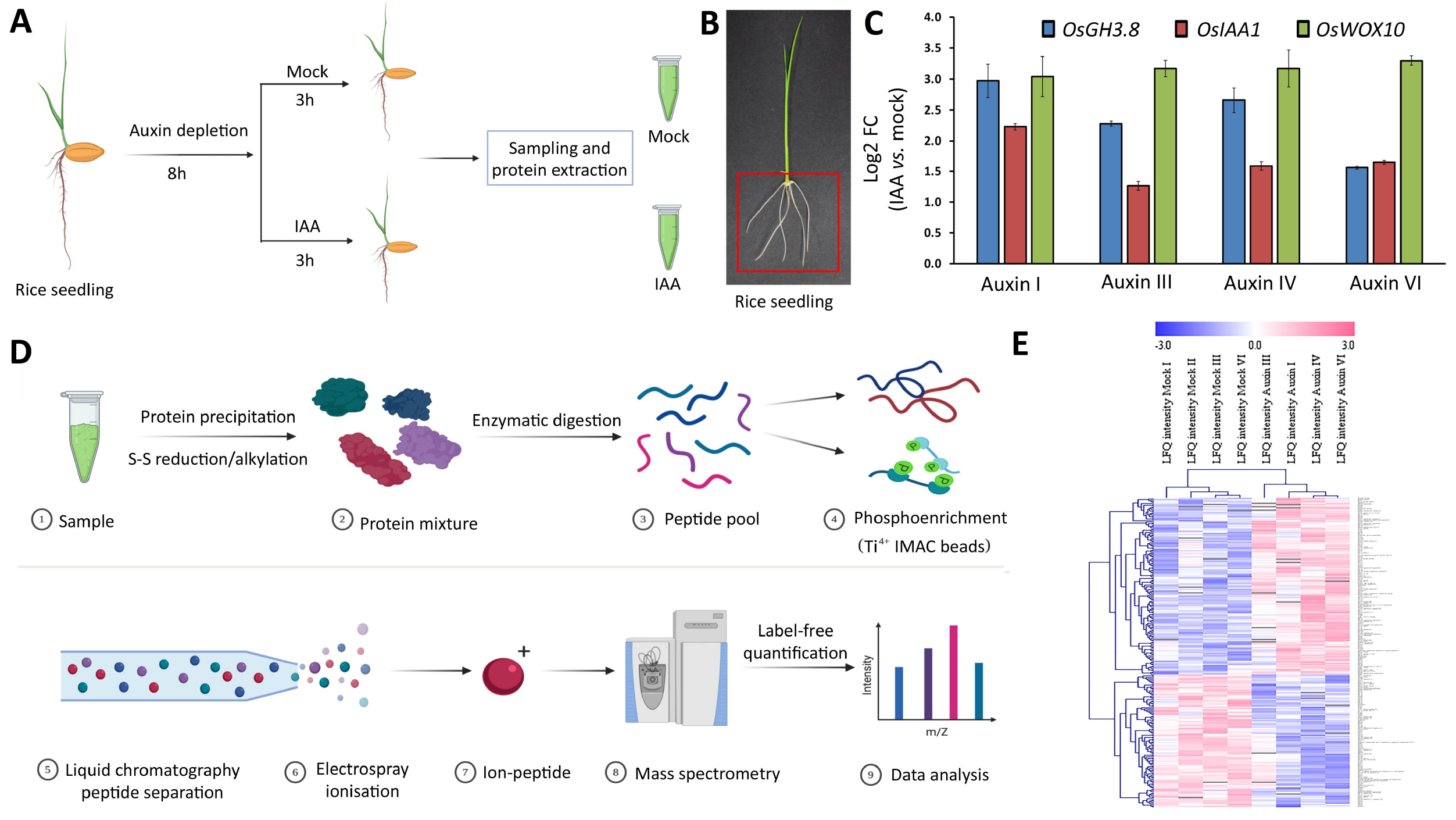
Workflow and differential (phospho)proteome response upon auxin induction. (A) Experimental workflow of plant treatment with control (mock) and auxin (IAA). (B) The red box highlights the plant region sampled. The 2-3 mm coleoptile base region along with entire crown root system was collected from 6^th^ day old seedling (primary roots and seeds were excluded). (C) Validation of auxin induction by analysing expression of a few auxin-responsive genes by quantitative real-time PCR (qRT-PCR). (D) Workflow for the characterization of proteins and phosphoproteins differentially regulated in response to auxin induction in rice coleoptile base and crown roots. (D) Differential proteome response upon auxin induction. Heat map showing average log2 values of MaxLFQ intensities of the differentially expressed proteomics intensities of the significantly regulated proteins with p < 0.05.

### Biological processes associated with chromatin conformational change and gene expression are de-regulated by auxin signaling

To establish an association between differential proteins and biological processes, we performed a gene ontology (GO) enrichment using PLAZA MONOCOT 4.5. (Van Bel *et al.*, 2018) The GO enrichment analysis highlighted induced expression of proteins putatively involved in DNA unwinding, DNA geometric changes, chromatin-associated histone H3 proteins, and developmental process of adaxial and abaxial patterning (Fig. 2A), suggesting a role for auxin signaling in regulating dynamic chromatin conformational changes associated with DNA replication and gene expression during CR formation which are required for cell cycle re-activation during CRP initiation. In contrast, GO analysis of proteins whose abundance is decreased in response to auxin signaling showed the enrichment of cytokinin-responsive, ATPase-related proteins and abiotic and biotic stress signaling (Fig. 2B). These observations are also supported by associated molecular functions and cellular localization of proteins regulated by auxin-signaling (Supplementary Fig. S2; S3).

**Fig. 2.**
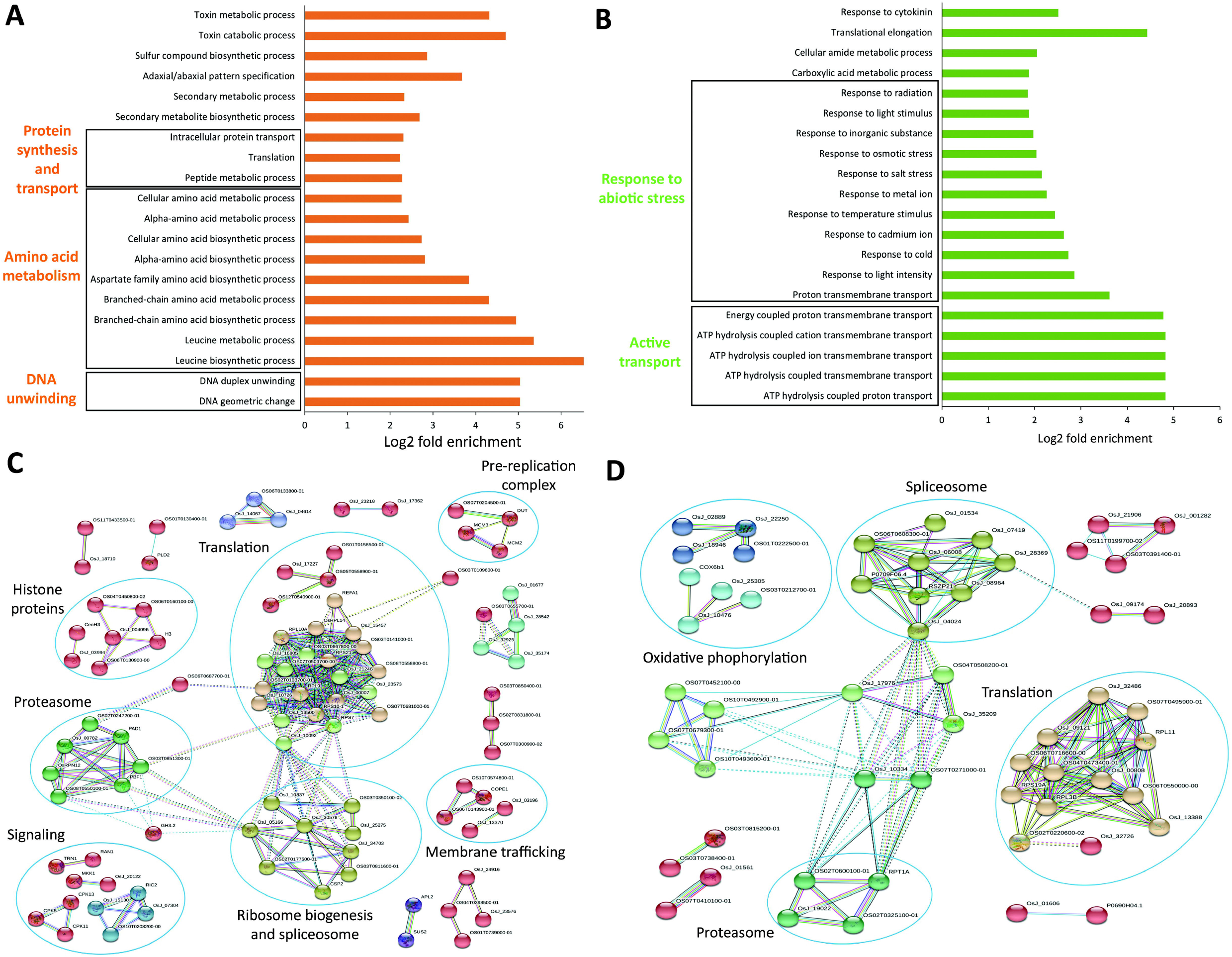
Gene ontology enrichment and protein interaction network analysis of differentially abundant proteins in response to auxin signaling. (A, B) Fold enrichment of top 20 GO terms associated with (A) upregulated proteins and (B) down-regulated proteins for biological processes. (C, D) Protein-protein interaction analysis of the auxin-induced (C) and auxin-repressed (D) proteins. STRING analysis was performed for the proteins showing differential abundance upon auxin induction. The disconnected nodes were removed from the network and the confidence score was set as > 0.9.

Next, to gain in-depth insight into the regulatory network functioning downstream of auxin signaling, a STRING network was constituted to decipher higher-level connections among differentially quantified proteins, including direct physical associations and indirect functional interactions (Szklarczyk *et al.*, 2019). STRING analysis showed a high degree of connectivity (Fig. 2C, D), suggesting a biological correlation among differentially regulated proteins. Both induced and repressed proteomes had significantly more interactions than expected; 272 edges compared to 199 expected and 149 edges compared to 131 expected, for induced and repressed proteins, respectively (Fig. 2C, D). Interestingly, the central clusters observed in both datasets were dominated by components of spliceosome machinery, translation-related processes, and proteasome. All these observations together suggest that auxin signaling also regulates post-transcriptional, translational and post-translational gene regulation programs.

### Protein abundance of cell signaling and cell cycle genes are regulated by auxin signaling

We also observed that proteins involved in endocytosis and trafficking and signal transduction were upregulated upon auxin treatment (Supplementary Dataset S1), which is consistent with the fact that the protein clusters especially present in the auxin-induced protein networks include membrane trafficking and signaling-related proteins (Fig. 2C). For example, *Osksr7* (Os11g24560), which encodes for the SEC23 subunit of the coat protein complex II (COPII) involved in ER-to-Golgi transport, a MAR binding filament protein *MFP1* (Os03g11060), and a calcium/calmodulin-dependent protein kinase *CAMK_like.27* (Os04g49510), displayed increased protein abundance in response to auxin signaling (Fig. 3A-C). On the other hand, OsMPK1 (Os06g06090), which belongs to plant mitogen-activated protein kinase family, and OsPRP1, a novel proline-rich glycoprotein, were down-regulated in response to auxin treatment (Fig. 3D, E).

**Fig. 3.**
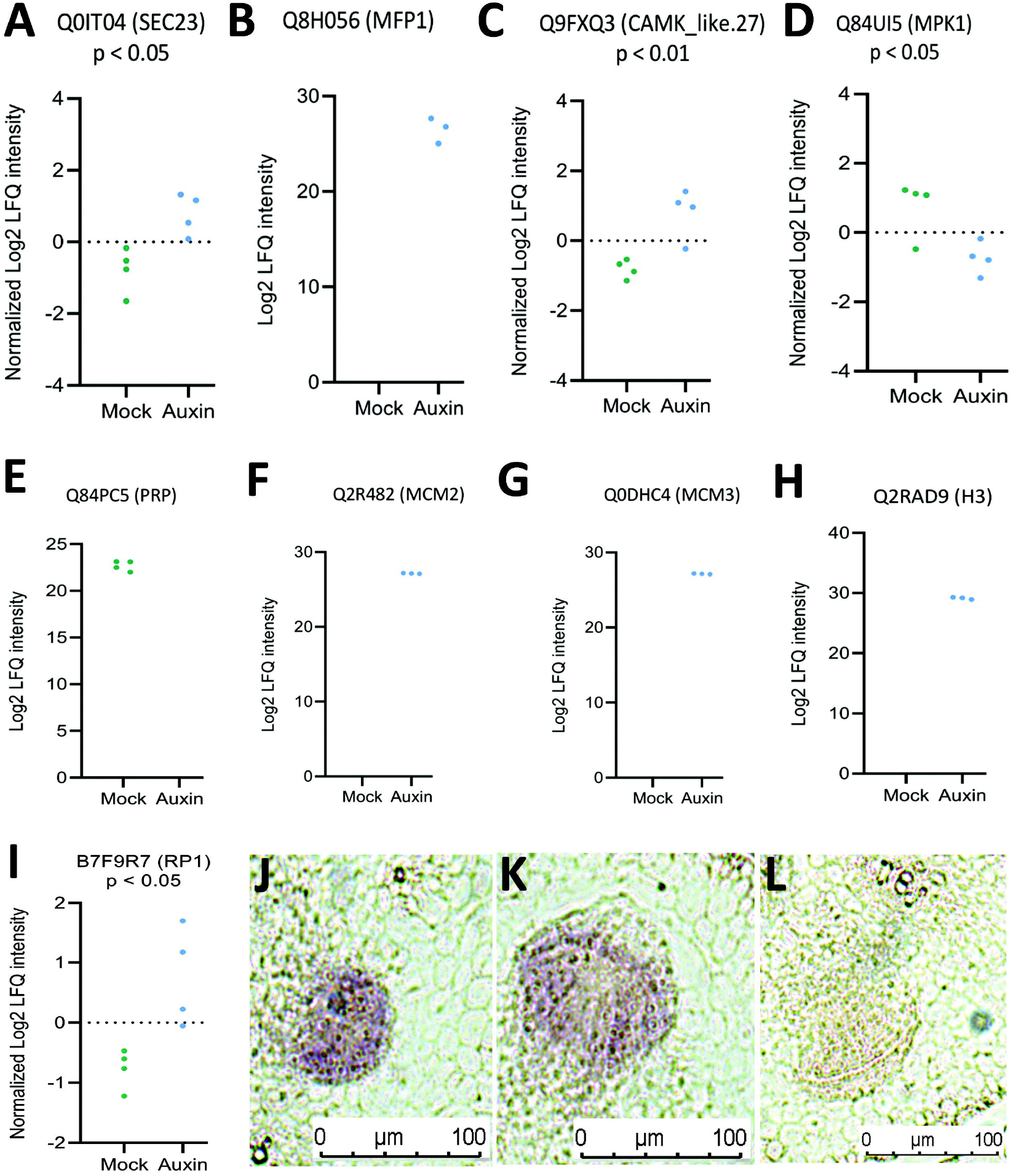
Auxin regulates proteins associated with gene expression, cell-signaling and cell cycle. (A-E) Graphs showing differential protein levels for the candidate proteins associated with cell signaling. (A) *Osksr7* (Os11g24560) encoding for the SEC23 subunit of the coat protein complex II (COPII), (B) a MAR binding filament protein MFP1 (Os03g11060), (C) a calcium/calmodulin-dependent protein kinase OsCAMK_like.27 (Os04g49510), (D) a mitogen-activated protein kinases family protein OsMPK1, and (E) a proline-rich glycoprotein OsPRP. (E-G) Proteins associated with DNA replication and maintenance. (F, G) minichromosome maintenance proteins OsMCM2 (Os11g29380) and OsMCM3 (Os05g39850) and (H) histone H3 protein. (I) Graphs showing differential protein levels for the protein encoding ribosomal protein RP1. (J-L) Tissue specific expression pattern of *RP1* gene during CR development, hybridized with (J, K) anti-sense DIG-RNA probe and (L) sense probe. Bars: (H-J) 100 mm.

CRP establishment requires re-activation of cell cycle and cell division in the ground meristem tissues in response to endogenous cues. Strikingly, we observed upregulation of proteins associated with DNA replication during the cell cycle, such as histone proteins, pre-replication complex, and minichromosome maintenance proteins, MCM2 (Os11g29380) and MCM3 (Os05g39850) (Fig. 3F-H). Thus, auxin signaling activates cellular signaling required for re-entry of differentiated cells into cell cycle program to initiate CRP.

### Spatial gene expression pattern of auxin-regulated proteins during CR development

Next, to validate the involvement of proteins whose abundance was regulated by auxin, in CR development, we analysed their spatial transcript distribution patterns during the initiation of CRP and its outgrowth using our recently generated genome-wide high-resolution expression dataset (Supplementary Fig. S4A; Garg *et al.*, 2020). Out of 201 proteins whose abundance was increased, 18 corresponding genes including ethylene biosynthesis enzymes, 1-aminocyclopropane-1-carboxylate oxidases (OsACO1 and OsACO2), two tRNA synthetases, a regulatory subunit of serine/threonine-protein phosphatase, components of proteasome machinery, and histone H3 were exclusively induced during CRP initiation, whereas seven genes were explicitly induced during CRP outgrowth (Supplementary Fig. S5B, C; Supplementary Dataset S2). However, the expression level of 64 genes, mainly including proteins involved in various steps of protein biosynthesis and degradation, and cell signaling, was commonly induced during CRP initiation and outgrowth (Supplementary Fig. S5D; Supplementary Dataset S2). The expression of 27 genes was higher during CRP initiation and their expression was reduced when CRP progress to the outgrowth stage and 27 genes displayed the opposite pattern (Supplementary Fig. S5E; Supplementary Dataset S2). A similar analysis for 133 proteins whose abundance was decreased upon IAA treatment identified genes with dynamic expression pattern during CRP initiation and outgrowth (Supplementary Fig. S5A-E; Supplementary Dataset S2). These genes were also analysed for their expression pattern in different root zones using the CoNekT database (Proost and Mutwil, 2018). Strikingly, 40.7% genes of all the differentially regulated proteins, were abundantly expressed in the root meristematic zone (Supplementary Fig. S5F). All these data together show that auxin-regulated genes display a dynamic expression pattern during CRP development and most of them might have a function in the root meristem.

Further, to validate spatial activation of the biological process during CRP development, we analysed the expression pattern of a ribosomal protein of L1P family, Os01g64090, involved in the translation-related process. The protein abundance of Os01g64090 is increased upon auxin treatment (Fig. 3I) and it is transcriptionally activated during CRP initiation and outgrowth (Supplementary Dataset S2). RNA *in situ* hybridization analysis using anti-sense RNA probes against Os01g64090 further confirmed that transcription of ribosomal proteins is specifically and strongly activated in the developing CRP (Fig. 3J, K). Its expression was induced during CRP initiation (Fig. 3J) and continued during CRP outgrowth (Fig. 3K). We did not observe any signal, when probed with sense riboprobes for the gene (Fig. 3L).

### Auxin signaling regulates amino acid metabolism

In GO analysis, we also observed an over-representation of GO terms associated with cellular alpha amino acid biosynthesis and metabolism, protein synthesis and protein transport in the proteins upregulated by auxin signaling (Fig. 2A). The amino acid biosynthesis and metabolism category was primarily dominated by aspartate family, and branched-chain amino acid (BCAA) that includes valine, leucine and isoleucine. In addition to being essential components of protein translation, amino acids also act as a signal to regulate other processes including gene expression and metabolic activities (Meijer and Dubbelhuis, 2004; Kimball and Jefferson, 2006).

To further confirm effect of auxin treatment on amino acid metabolism, we performed GC-MS based amino acid profiling of rice coleoptile base and emerged CRs in response to auxin treatment. The supervised partial least squares discriminant analysis (PLS-DA) and the 2D score plots obtained from the PLS-DA test showed that the amino acids from mock and IAA treated samples did not overlap with each other, indicating an altered state of metabolite levels (Fig. 4A). The first two components (component 1 and 2) of PLS-DA accounted for 41.9 % and 21.6 % variance among samples, respectively. This analysis identified than abundance of 12 amino acids including a few members of aspartate family such as aspartic acid, lysine, threonine and isoleucine was altered upon auxin treatment (Fig. 4B, Supplementary Fig. S5A-C; Supplementary Table S1). The level of all three BCAAs, i.e. valine, leucine and isoleucine was increased upon induction of auxin signaling during CR development in rice (Fig. 4B, Supplementary Fig. S5A-C; Supplementary Table S1). Next, the influence of the amino acids (independent variable) in determining the dependent variable (effect of auxin treatment) was estimated by VIP scores where amino acids with VIP score (≥1) were considered as important marker to play crucial roles in distinguishing auxin treated samples. Eight differential amino acids, L-phenylalanine L-lysine, L-5-oxoproline, L-valine, isoleucine, L-alanine, L- threonine, and L-leucine were detected with VIP >1 (Fig. 4C). Thus, both proteome and metabolite profiling demonstrate regulation of amino acid metabolic flux by auxin signaling during CR development.

**Fig. 4.**
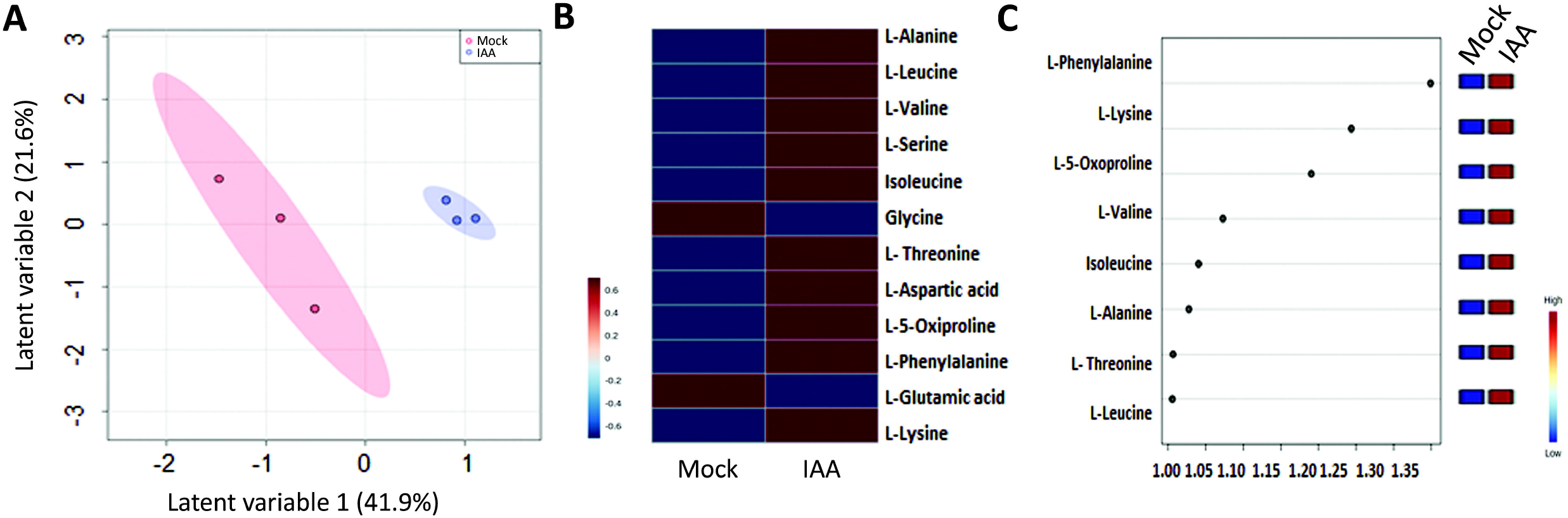
Metabolic profiling of rice coleoptile base and emerged CRs in response to auxin treatment. (A-C) Multivariate statistical analysis for PLS-DA 2D scores plot (A), heatmap generated by Metaboanalyst 4.0 (B), and important features, PLS-DA, VIP scores, (C) of differentially accumulating amino acids detected by GC-MS from mock and IAA treated samples. In scores plot, amino acids from IAA treated and mock samples didn’t overlap with each other, indicating an altered state of amino acid levels.

### Auxin signaling regulates its homeostasis during adventitious root development

The active pool of endogenous auxin is crucial for controlled activation of auxin signaling which is maintained by a balance between auxin biosynthesis and degradation/inactivation (Peer, 2013). YUCCA genes are flavin-containing monooxygenases (FMOs), involved in auxin biosynthesis, whereas members of the GH3 gene family inactivate the free intracellular auxin pools by conjugating them with other biomolecules in plants (Staswick *et al.*, 2005; Schlaich, 2007; Yadav *et al.*, 2011). We observed that the protein level of an *OsFMO* (Os07g02100) gene, and a GH3 family member, *OsGH3-2* (Os01g55940) were induced upon auxin treatment (Fig. 5A, B). The expression of *OsGH3-2* is also transcriptionally induced by auxin signaling (Neogy *et al.*, 2019). The expression of *OsGH3-2* was sharply activated during CRP initiation and outgrowth, whereas Os07g02100 has higher expression during CRP initiation which is reduced during CRP outgrowth (Supplementary Dataset S2). The dynamic expression pattern of Os07g02100 was consistent during CRP development as also validated by RNA *in situ* hybridization. It was strongly activated during CRP initiation (Fig. 5C-F) and continued to express during CRP outgrowth (Fig. 5F). However, the sense probe did not give any signal in developing CRP (Fig. 5G).

**Fig. 5.**
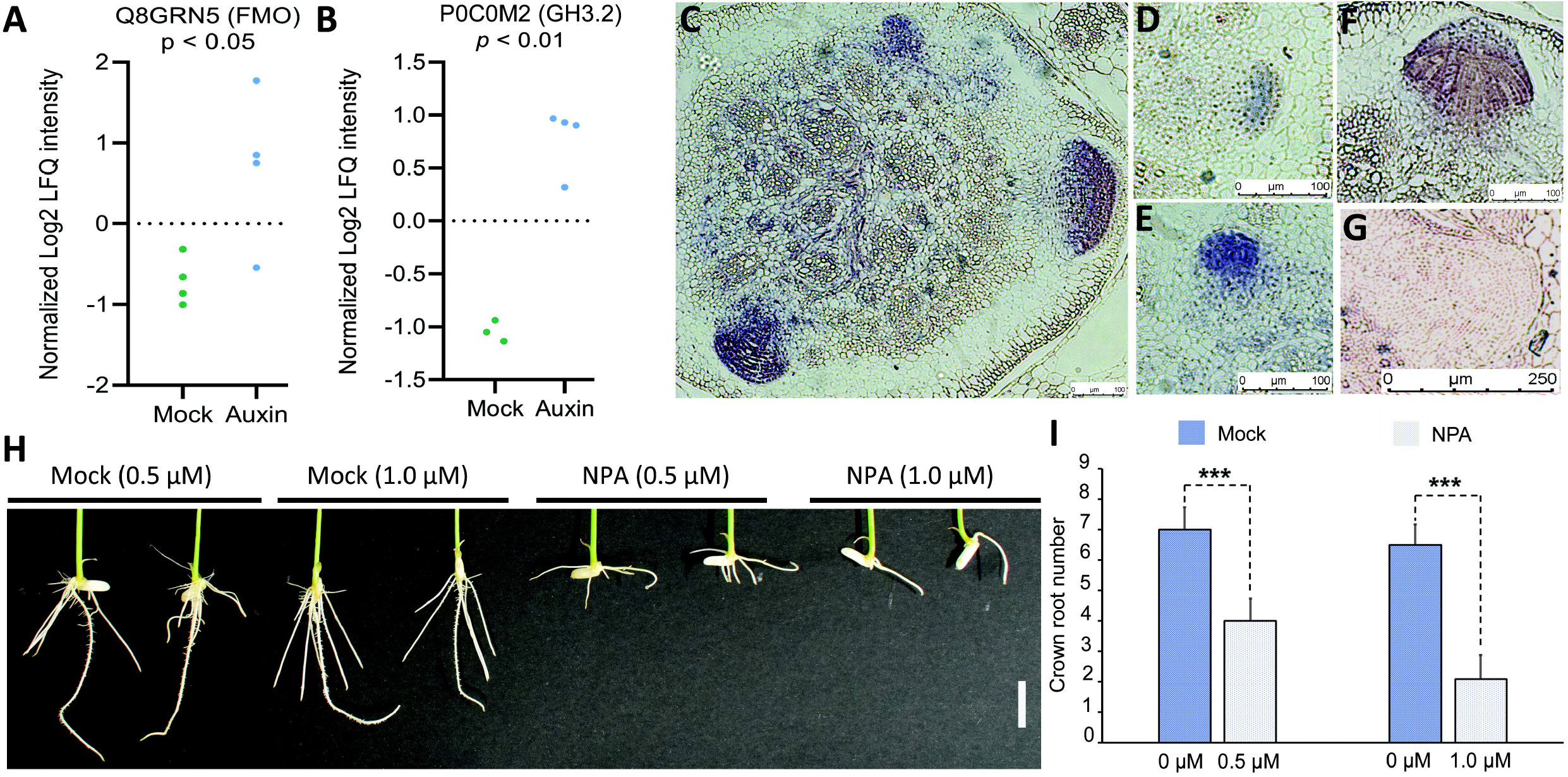
Auxin signaling during crown root development. (A, B) Graphs showing differential protein levels for the flavin-containing monooxygenases FMO, Os07g02100 (A) and a GH3 family member, *OsGH3-2* (Os01g55940) proteins (B). (C-G) Tissue specific expression pattern of *OsFMO* gene during CR development, hybridized with anti-sense DIG-RNA probe (C-F) and sense probe (G). (H) Root architecture phenotypic of 6^th^ day old seedling treated with auxin signaling inhibitor, the naphthylphthalamic acid (NPA). (I) Number of crown roots was reduced in NPA treated plants. (***, p<0.0001), Bars: (C)-(F) 100 μm, (G) 250μm, (H) 1cm.

To further functionally validate this observation, we interfered with auxin signaling using the pharmacological inhibitor of auxin transport, naphthylphthalamic acid (NPA). NPA treatment resulted in a significant decrease in the number and growth of CRs in rice (Fig. 5H-I). Thus, translational regulation of FMO and GH3 by auxin signaling and their temporal activation in the developing CRP are also corroborated with their function in rice CR development as alteration of auxin homeostasis either by NPA treatment or over-expression of *OsGH3-2* resulting in reduced CR formation (Du *et al.*, 2012).

### Auxin-mediated activation of ethylene signaling is required for crown root initiation

An interaction between auxin and ethylene signaling is known to regulate adventitious root development (Veloccia *et al.*, 2016). We observed that protein abundance of ethylene biosynthesis enzymes, 1-aminocyclopropane-1-carboxylate oxidase, OsACO1 (Os03g04410) and OsACO2 (Os09g27750) was induced upon auxin treatment (Fig. 6A; Supplementary Dataset S1). Further, the protein level of an ethylene inducible acireductone dioxygenase enzyme OsARD1 (Os10g28350) and of PDX11 (Os07g01020) was oppositely altered upon auxin treatment (Fig. 6B, C). The expression of *OsACO* genes is also induced by auxin at the transcription level (Neogy *et al.*, 2019) and its transcription is exclusively and transiently activated during CRP initiation (Supplementary Dataset S2), suggesting that it may be functioning during CRP initiation. To further validate activation of ethylene biosynthesis during CRP establishment, temporal and spatial expression of *OsACO1* was analysed. *OsACO1* was exclusively activated in the CRP (Fig. 6D) and its onset of expression occurred during CRP establishment (Fig. 6E). The expression continued during CRP outgrowth (Fig. 6F) and emergence (Fig. 6G, H).

**Fig. 6.**
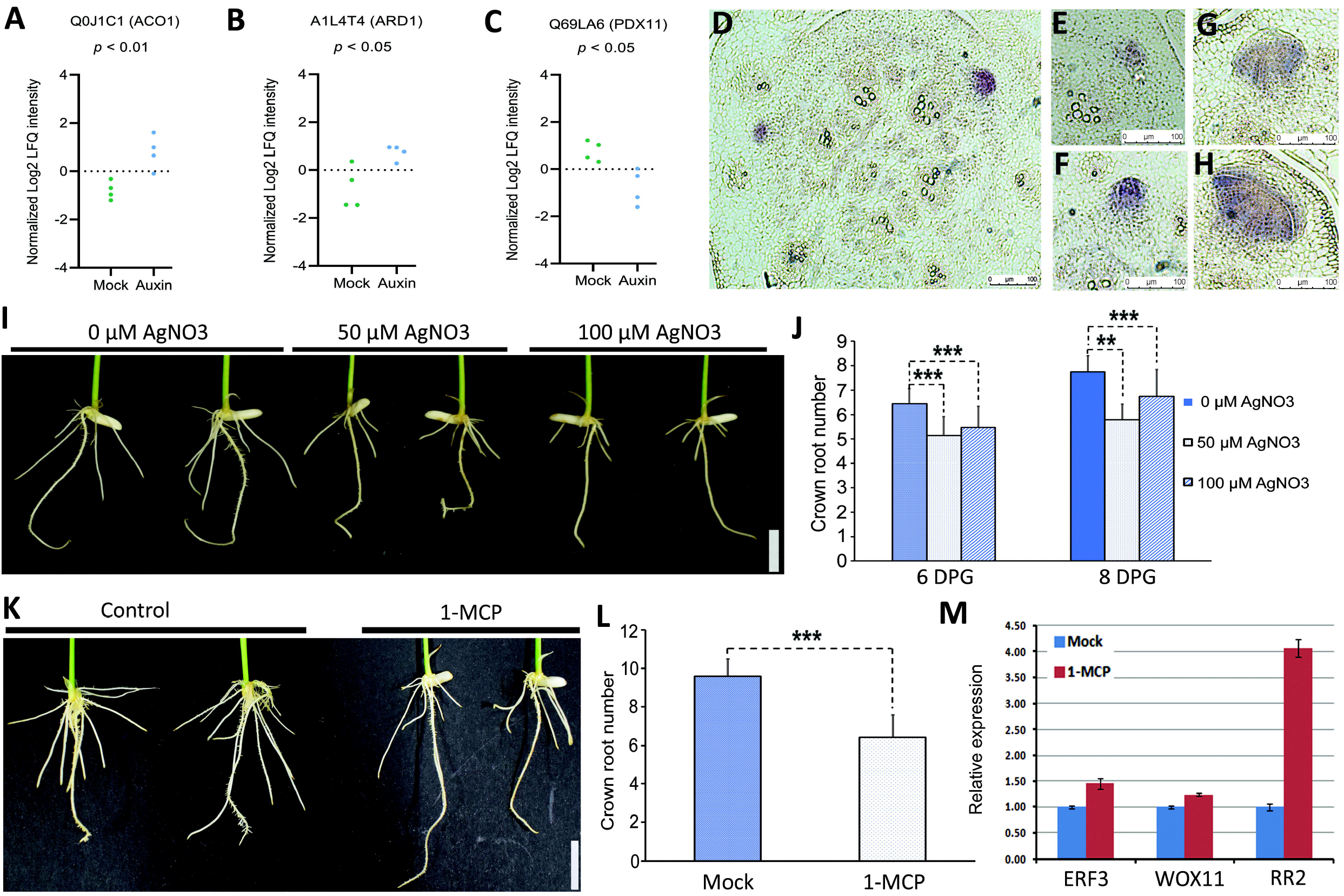
Auxin activates ethylene signaling during crown root formation. (A-C) Graphs showing differential protein levels of an ethylene biosynthesis enzyme, the aminocyclopropane-1-carboxylate oxidases encoding *OsACO1* (A), and ethylene inducible proteins, an acireductone dioxygenase enzyme *OsARD1* (B), and a putative SOR/SNZ family protein, *OsPDX11* (C). (D-H) Temporal-spatial expression pattern of *OsACO1* gene during CR development. (I) Root architecture phenotype of rice seedling upon silver nitrate (AgNO_3_) treatment. (J) Quantitative representation of crown root number in rice seedling treated with AgNO_3_. (K) Root architecture of rice seedling treated with 1-Methylcyclopropene (1-MCP), a competitive inhibitor of ethylene signaling. (L) Number of crown roots was significantly reduced in the seedling inhibited for ethylene signaling using 1-MCP. (M) Gene expression analysis of a key regulatory module (ERF3-WOX11-RR2) of crown root development using q-RT PCR. Ethylene signaling repressed expression of cytokinin response regulator, RR2, a negative regulator of crown root emergence. (DPG, day post germination), (**, p < 0.01; ***, p < 0.0001). Bars: (D)-(H) 100 μm, (I), (K) 1cm.

Next, to functionally confirm that ethylene signaling is involved in rice CR development, we studied consequences of ethylene signaling inhibition on CR formation. Silver ions, Ag(I) effectively block ethylene action in plants (Beyer, 1976), and therefore we germinated rice seeds in the presence of different concentrations of silver nitrate (AgNO_3_). We observed that root architecture was altered and the CR number was significantly reduced upon AgNO_3_ treatment (Fig. 6I, J). Furthermore, we also used 1-Methylcyclopropene (1-MCP), a competitive inhibitor of ethylene signaling by tightly binding with the ethylene receptor (Sisler *et al.*, 1996; Sisler and Blankenship, 1996). We observed similar effects on root architecture and CR number with 1-MCP treatment (Fig. 6K, L). It is important to note that the effects of these inhibitors were more prominent on CRs and that only a marginal or no effect was observed on primary roots. Expression analysis of genes of a key regulatory module (ERF3-WOX11-RR2) of CR development showed that the expression of *OsRR2* but not *ERF3* and *WOX11*, was induced upon 1-MCP treatment (Fig. 6M). All these together suggest that auxin activates ethylene signaling specifically in the CRP that regulates rice CR development through a cytokinin response regulator.

### Global (phospho)proteome analysis reveals that WOX11, a key regulator of CR development is phosphorylated in rice

Reversible protein phosphorylation, a critical post-translational protein modification, provides a regulatory switch for controlling protein activity, sub-cellular localization, and molecular interactions during various biological processes including growth and development (Cohen, 2002; Yadav *et al.*, 2020). Therefore, to reveal phosphorylation-mediated post-translational regulation during rice CR development, we performed LC-MS/MS-based global phosphoproteome analyses of the rice coleoptile base and emerged crown roots using a previously optimized protocol for *Arabidopsis thaliana* and maize (Vu *et al.*, 2016). A total of 8220 phosphosites that were identified through Ti^4+^ IMAC enrichment analysis could be mapped to 1594 phosphoproteins (Supplementary Dataset S3). An average Pearson correlation of 0.849 and 0.859 for mock and auxin-treated samples, respectively showed quantitative reproducibility of replicates (Supplementary Fig. S6). In the total identified phosphosites, the contribution of pSer is 90%, pThr is 9.4% and of pTyr is 0.7%. (Supplementary Fig. S7), which is similar to past studies from different tissues of monocots and dicots (Nakagami *et al.*, 2010; Wang *et al.*, 2017). Since tyrosine phosphorylation is very limited in the plant kingdom, of the total 8220 phosphosites, only 54 were phosphorylated at a tyrosine residue.

Members of the WUSCHEL-related homeobox (WOX) transcription factor family play a key role in regulating root development. We observed phosphorylation of rice WOX11 in CRs (Supplementary Dataset S3). WOX11 plays a crucial role during rice CR development and it directly interacts with ERF3 to represses the expression of the cytokinin response regulator *RR2* during the emergence of crown roots (Zhao *et al.*, 2009, 2015). We, therefore, speculate that phosphorylation/dephosphorylation of WOX11 might affect its interaction and/or activities during CR development.

### Auxin-triggered differential phosphoproteome during CR development

Next, we also performed LC-MS/MS-based auxin-dependent phosphoproteome analyses of the rice coleoptile base and emerged crown roots. Previous differential phosphoproteome studies of rice tissues lacked normalization to the protein abundance. As protein levels for many phosphosites could not be deduced, because no non-phosphorylated peptides of the corresponding proteins were detected, we first subjected the non-normalized phosphosites dataset to a two-sample test (p < 0.05). Based on the phosphoproteome data alone, we identified 42 phosphosites which were significantly more phosphorylated, whereas 24 phosphosites were significantly less phosphorylated upon IAA treatment (Fig. 7A; Supplementary Dataset S3). More importantly, of these, 25 phosphosites were uniquely detected upon IAA treatment but not in mock-treated samples and 10 phosphosites were uniquely detected only in the mock-treated samples (Supplementary Dataset S3).

**Fig. 7.**
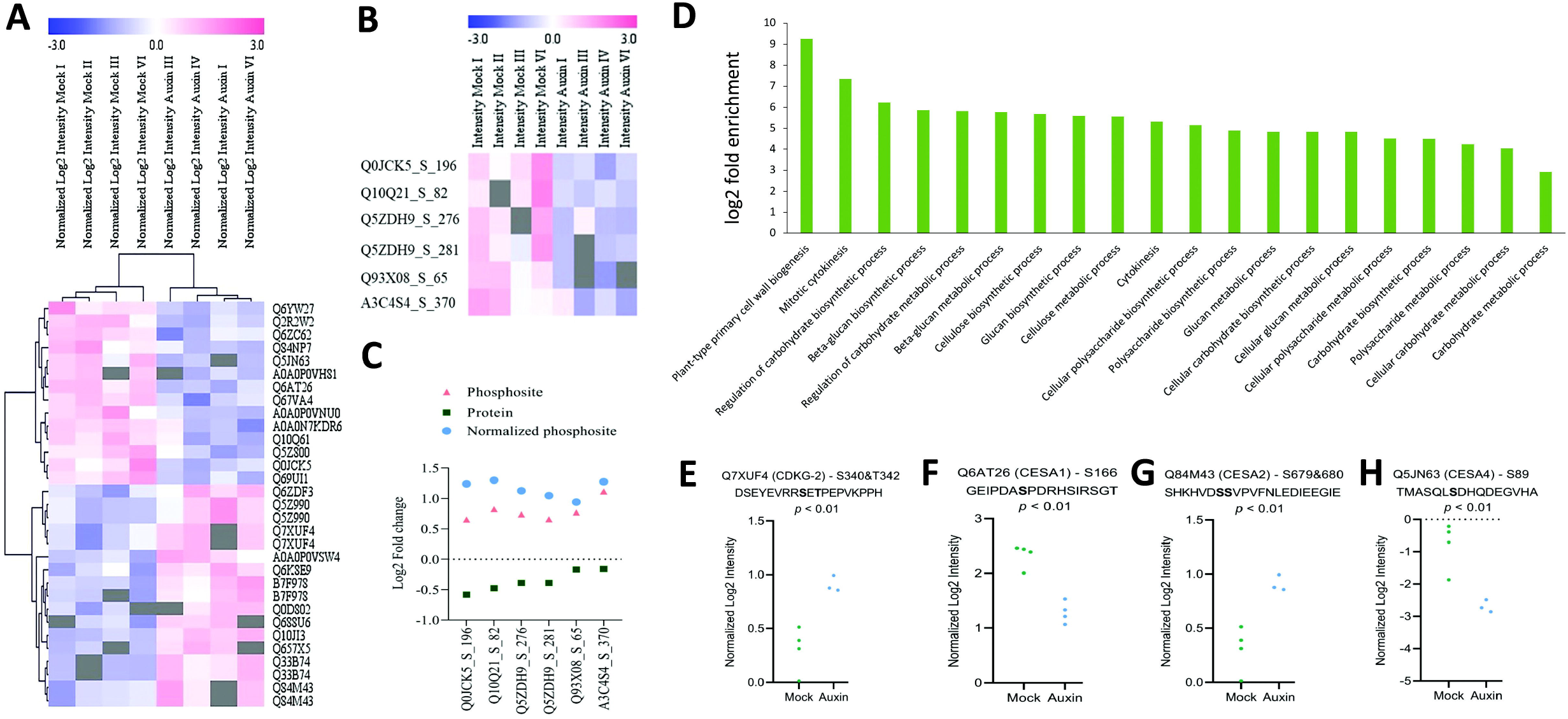
Differential phosphoproteome response upon auxin induction. (A) Heatmap showing average Log2 transformed intensities of the phosphosites for the significantly regulated phosphorylated peptides with p < 0.05. (B, C) A normalized auxin-triggered phosphoproteome. Significantly de-regulated phosphopeptides were normalized by subtracting the log2 fold change of the protein abundance from the log2 fold change of the phosphopeptide. (D) Cell division and cell wall associated biological process GO terms were enriched in the proteins differentially phosphorylated upon auxin treatment. The analysis was performed using PLAZA monocot 4.5 (p-value < 0.05). Graphical representation of differential phosphorylation of (E) a cell cycle regulating kinase (CDKG-2) and cellulose biosynthesis proteins (F) CESA1, (G) CESA2 and (H) CESA4.

Taking into account that the outcome of differential phosphorylation data can be influenced by the difference in protein levels, we further normalized the intensities of the phosphopeptides to the protein intensities (Vu *et al.*, 2016). We mapped the significantly de-regulated phosphopeptides for which the proteins were not detected among the differential proteins (Supplementary Figure S8), and whole phosphoproteome, on the whole proteome data to identify phosphorylation events that were not identified as significant. We identified 944 phosphosites (corresponding to 469 phosphorylated proteins) that could be normalized for their protein abundance. Of those, six phosphosites were differentially regulated by auxin signaling based on a two-sample test (p < 0.05) on the normalized phosphosites intensities (Fig. 7B, C; Supplementary Dataset S3). This small set of phosphorylation events that are fully due to kinase and phosphatase activity and not because of changes in protein abundance. This list includes a putative PB1 domain-containing protein (Q5ZDH9), a UTP-glucose-1-phosphate uridylyltransferase (or UDP–glucose pyrophosphorylase/UGPase) (Q93X08), a GDP-mannose 3,5-epimerase 1 (A3C4S4), a Ubiquitin binding domain (UBD) containing protein (Q0JCK5) and a mitochondrial peptidase (Q10Q21) (Supplementary Dataset S3).

### Auxin-dependent phosphorylation is required for cell cycle activation, cell signaling and cell wall synthesis during root organogenesis

Our in-depth GO enrichment analysis of the proteins revealed that a few cell cycle-associated genes were differentially phosphorylated in response to auxin signaling (Fig. 7D; Supplementary Dataset S3). Interestingly, we observed that a member of the rice CDK family, *CYCLIN-DEPENDENT KINASE G-2 (OsCDKG;2),* showed induced phosphorylation at two phosphosites, S 340 and T 342, upon auxin treatment (Fig. 7E). Importantly, *AtCDKG;2* regulates adventitious root development in *Arabidopsis* and calli derived from *cdkg;2* mutants, failed to induce roots (Zabicki *et al.*, 2013). Furthermore, we observed the phosphorylation of signaling-related proteins, ATPase and GTPases, and cell wall associated proteins (Supplementary Dataset S3). Auxin activates H^+^-ATPases and cell wall synthesis genes, leading to apoplastic acidification, which further triggers Ca^2+^ pumps. Finally, the Ca^+^ signal upregulates RHO OF PLANTS (ROP) GUANOSINE-5’-TRIPHOSPHATASES (GTPases) and promotes the delivery of new cell wall components (Fu *et al.*, 2002; Gu *et al.*, 2004). In addition to UGPase and GDP-mannose, we also observed differential phosphorylation of three rice *CESA* genes, *CESA1, CESA2* and *CESA4* (Fig. 7F-H; Supplementary Fig. S9; Supplementary Dataset S3).

Next, we analysed spatial RNA expression pattern of the genes displaying differential phosphorylation in response to auxin signaling, during CRP initiation and outgrowth, and different zones of growing CR. Among the proteins which were phosphorylated upon IAA treatment, the expression of relatively larger number of genes were decreased in developing CRP as compared to control competent tissues, whereas the pattern is opposite for the proteins dephosphorylated upon auxin signaling (Supplementary Fig. 10A-D; Supplementary Dataset S4). The transcript distribution analysis of auxin-regulated differentially phosphorylated proteins in different root zones of growing crown roots revealed that 54.2% of total genes were abundantly expressed in the differentiation zone of growing roots (Supplementary Fig. 10E). Therefore, suggesting the importance of auxin signaling during the process of cell differentiation.

## Discussion

Adventitious/crown root formation initiates with the specification of crown root founder cells in the innermost ground tissues, peripheral to the vascular tissues, of rice coleoptile/stem base in response to positional cues (Itoh *et al.*, 2005). The founder cells re-enter in the cell cycle and a set of formative cell divisions results in the formation of three layers of fundamental tissues in a CRP. During the later stage, root cell differentiation and tissue patterning follows CRP outgrowth which eventually emerges out as CR (Itoh *et al.*, 2005). Our study (phopho) proteome and metabolome analysis reveals that auxin regulates a plethora of proteins and metabolites belonging to diverse functional categories associated with gene expression, cell cycle, ethylene signaling and cell wall synthesis during crown root formation in rice.

### Auxin-regulated gene expression re-activates cell cycle in CRP founder cells

Cell cycle re-activation will require expression of genes required for DNA replication. A correlation exists between auxin signaling and chromatin accessibility, which is associated with DNA replication and gene expression (Hasegawa *et al.*, 2018). In this study, we observed changes in abundance of proteins involved in the process of DNA unwinding, DNA geometric changes, RNA splicing, and amino acid and protein metabolism. Further, the proteins related to amino acid metabolism, protein synthesis and transport were upregulated by auxin signaling. In plants, amino acids not only play a vital role as building blocks of protein translation, but also act as a signal to regulate gene expression, translation and metabolic activities (Meijer and Dubbelhuis, 2004; Kimball and Jefferson, 2006). One of the BCAA mutants, *low isoleucine biosynthesis (lib),* which is partially deficient in isoleucine, displays defects in cell proliferation and expansion during root development (Yu *et al.*, 2013). It is reported that coordinated actions of auxin, Rho-like small GTPases (ROPs) and target of rapamycin (TOR) signaling are involved in the translation re-initiation from mRNAs (Schepetilnikov and Ryabova, 2017). Moreover, auxin-inducible genes, *SAUR62* and *SAUR75* regulate ribosome abundance and assembly (He *et al.*, 2018). These results suggest a role for auxin signaling in regulating dynamic chromatin conformational changes associated with DNA replication and gene expression at the translational and/or post-translational level during crown root formation. Also, the induced levels of histone, pre-replication complex and signaling proteins (Calcium-dependent protein kinases) might serve as the initial stage preparation for the cell cycle initiation and cell division.

An auxin maximum is a prerequisite for a cell to divide and it also controls reversible modification of key regulators of cell cycle (Braun *et al.*, 2008; del Pozo and Manzano, 2014). Cyclin-dependent kinases (CDKs) are ser/thr protein kinases and play a key role in regulating cell cycle in eukaryotes (Lees, 1995; Morgan, 1997). CDK*-*activating kinases (CAKs) phosphorylate at the threonine site in the T-loop of CDKs to activate them (Shimotohno *et al.*, 2003). OsCDKG;2 is differentially phosphorylated in response to auxin signaling. The phylogenetic analysis of CDKs shows that OsCDKG;2 belongs to a subclade consisting of OsCDKG;1, AtCDKG;1, and AtCDKG;2 (Guo *et al.*, 2007). AtCDKG;2 is shown to be involved in regulating *Arabidopsis* adventitious root development (Zabicki *et al.*, 2013). In addition to the role of AtCKDG;1 in the cell cycle, it has also been shown to be associated with alternative splicing of *CALLOSE SYNTHASE5* (*CalS5*), a regulator of symplastic cell-cell communication and pollen cell wall formation (Huang *et al.*, 2013). The expression of *OsCDKG;1* and *OsCDKG;2* was repressed by cytokinin signaling (Guo *et al.*, 2007), corroborating with their putative positive function during CRP initiation. We speculate that auxin-dependent phosphorylation of OsCDKG;2 might be amongst the early signaling events in re-activating the cell cycle in the competent cells.

### Cellular and auxin signaling is translationally regulated during CRP development

Our study shows that some cell signaling including MAP kinase is translationally regulated by auxin. An earlier study in *Arabidopsis* reported that MAP kinase signaling pathways are induced upon auxin treatment and interact with auxin-signaling pathway and generate a feed-back regulatory loop (Mockaitis and Howell, 2000). Further, in rice, OsMPK1 interacts with OsAux/LAX1 protein and MAP kinase cascade is involved in auxin signaling events (Mohanta *et al.*, 2015). *OsPRP1* regulates root growth (Akiyama and Pillai, 2003; Tseng *et al.*, 2013) and interacts with cell-wall related proteins and suppresses cell expansion, suggesting the involvement of *OsPRP1* in auxin-mediated regulation of cell expansion.

The auxin-induced protein network also highlights the abundance of ribosomal proteins, membrane trafficking and cell-signaling processes. Auxin have been known to regulate the transcription and translation pattern of ribosomal proteins (Gantt and Key, 1985; Beltrán-Peña *et al.*, 2002). Further, lipid-based metabolic processes are known to modulate auxin-mediated endomembrane trafficking pathways and tissue-differentiation downstream of ribosomal proteins, which implicates the role of ribosomal proteins as a translational regulator in response to auxin (Li *et al.*, 2015*b*). In addition, ribosomal proteins also play an important role in various plant growth and developmental processes ( Byrne, 2009). In rice, NAL21 encodes a ribosomal small subunit protein RPS3A and the *nal1* mutant was found to be aberrant in auxin responses (Uzair *et al.*, 2021). The expression of *OsGH3-2* is also transcriptionally induced by auxin signaling (Neogy *et al.*, 2019). In our study, we observed induced protein levels of FMO and OsGH3-2 proteins. *GH3* genes regulate the free intracellular auxin levels by conjugating IAA with the amino acids in plants and alteration of their endogenous expression results in defects in root architecture (Staswick *et al.*, 2005; Yadav *et al.*, 2011). On the basis of previous studies and our data, we implicate that auxin induces various ribosomal proteins which in turn regulate the protein synthesis of various auxin-signaling components. Moreover, components of the proteasome machinery, which generates a feed-back loop regulatory module with auxin signaling, were also de-regulated upon auxin treatment. While 26S proteasome-mediated degradation of Aux/IAAs is required for the activation of auxin signaling, auxin signaling also activates PTRE1, a putative repressor of 26S proteasome (Yang *et al.*, 2016). This suggest that auxin regulates various cellular responses, which further tunes the expression of downstream auxin-signaling components.

### Auxin activates ethylene signaling during rice CR development

Ethylene signaling is known to induce adventitious root development in plants (Lorbiecke and Sauter, 1999; Verstraeten *et al.*, 2014; Lakehal and Bellini, 2019). ACO is involved in the rate-limiting step of ethylene biosynthesis (Houben and Van de Poel, 2019). During ethylene biosynthesis, Met is catalyzed by a two-step reactions to yield the ethylene precursor, the 1-aminocyclopropane-1-carboxylic acid (ACC) (Wang *et al.*, 2002). An ethylene inducible acireductone dioxygenase enzyme OsARD1 (Os10g28350) has a role in the methionine (Met) salvage pathway, thus, generates a feed-back regulatory module to regulate ethylene biosynthesis (Liang *et al.*, 2019). We show that auxin signaling activates ACO and OsARD1 protein levels, thus working upstream of ethylene signaling pathway. Among the down-regulated proteins, PDX11 (Os07g01020) encodes for a putative SOR/SNZ family protein and shares a close homology with the pyridoxine synthase gene AT5G01410 (Chen *et al.*, 2014). *Arabidopsis PDX1.3* plays a vital role during the post-embryonic shoot and root development, and also during abiotic stress conditions (Chen and Xiong, 2005). The *pdx1.3* vitamin B6 mutant is impaired in ethylene production, and reduced accumulation of auxin and SHORT-ROOT (SHR) transcription factor, which manifested root developmental defects (Boycheva *et al.*, 2015). All these observations together suggest that ethylene biosynthesis functions under the regulation of auxin signaling and might play a vital role during CR development.

Members of the WUSCHEL-related homeobox (WOX) transcription factor family play a key role in establishing and maintaining the stem cell niche in the root apical meristem (RAM). *Arabidopsis* WOX5, was shown to get phosphorylated by the ACR4 kinase at the Serine residue *in vitro* (Meyer *et al.*, 2015). In rice, a regulatory module comprising of ERF3-WOX11-RR2 play central role during CR initiation and emergence (Zhao *et al.*, 2015). Interaction of ERF3 and WOX11 proteins repress expression of cytokinin response regulator RR2 during CR emergence (Zhao *et al.*, 2015). We show that ethylene biosynthesis is activated by auxin that in turns repressed expression of RR2. However, we speculate that WOX11 activities may be regulated post-translationally, through phosphorylation.

### Cell wall related proteins are phosphorylated in response to auxin signaling

Interestingly, our list of phosphorylation events included proteins such as, UBD and mitochondrial peptidase, which might be involved in proteolysis process. UGPase functions in the synthesis of uridine diphosphoglucose (UDPG), which is a glucose donor for sugar metabolism and cell wall formation (Sandhoff *et al.*, 1992). Auxin is known to regulate cell-wall synthesis (Majda and Robert, 2018), which suggests that UGPase is one of the prime targets of auxin during cell-wall formation and post-translational modifications might regulate its activity. In addition to UGPase and GDP-mannose, we also observed differential phosphorylation of three rice CESA genes, CESA1, CESA2 and CESA4. Cellulose synthase (CESA) proteins play a central role in cell wall biosynthesis and post-translational modifications play an essential role in their regulation (Speicher *et al.*, 2018). CESA proteins have multiple phosphorylation sites residing in the N terminus and at the cytosolic loop. Several studies report the crucial role of CESA phosphorylation in maintaining anisotropic cell expansion in roots and hypocotyl and also partially through its interaction with the cellular microtubules. The *CESA4* tissue-specific expression pattern showed its abundance in elongating tissues, such as developing leaf blades, elongating internodes and roots (Tanaka *et al.*, 2003). Many shreds of evidence provide a link between cell wall function, ethylene and auxin, such as *fei1 fei2* double mutants that show reduced growth anisotropy due to reduced cellulose biosynthesis and the *fei1 fei2* phenotype is rescued by auxin biosynthesis genes (Xu *et al.*, 2008; Basu *et al.*, 2016). However, there remains a gap regarding the kinases that phosphorylate multiple residues of CESAs during various developmental processes.

In summary, our study shows that auxin regulates a plethora of proteins belonging to diverse functional categories, influencing the cellular proteome in a tissue-specific context, during CR formation in rice. The data provides a rich source of fore-mining novel protein functions. In particular, the peptides related to cell cycle, protein synthesis, protein localization and post-translational modifications shows abundance in response to auxin induction. Additionally, the differential phospho-proteome data highlights the processes related to cell-wall synthesis and cell division in a root-specific fashion, which makes them excellent candidates for future functional and regulation studies during trans-differentiation.

## Abbreviations

CR: crown root
CRP: crown root primordia
IAA: Indole-3-acetic acid
LC-MS: liquid chromatography–mass spectrometry
GC-MS: gas chromatography–mass spectrometry
DIG-UTP: Digoxigenin-11-Uridine triphosphate
qRT-PCR: quantitative real-time PCR

## Supplementary data

**Supplementary Fig. S1:** Pearson correlation coefficient for rice proteome data.

**Supplementary Fig. S2:** Gene ontology enrichment analysis performed for differentially regulated proteins upon auxin treatment. Classification of up-regulated proteins on the basis of (A) molecular function. Classification of down-regulated proteins on the basis of (B) molecular function.

**Supplementary Fig. S3:** Gene ontology enrichment analysis performed for differentially regulated proteins upon auxin treatment. Classification of up-regulated proteins on the basis of (A) cellular localization. Classification of down-regulated proteins on the basis of (B) cellular localization.

**Supplementary Fig. S4:** Differentially regulated proteins manifest dynamic RNA expression pattern during CRP development. (A) A schematic representation of crown root primordium (CRP) initiation, CRP outgrowth, CRP emergence and different root zones. (B-E) Heatmap showing RNA expression pattern of differentially regulated proteins during CRP initiation and CRP outgrowth. (B, C) Genes specifically de-regulated during CRP initiation (B) or during CRP outgrowth (C), with respect to control tissues. (D) Genes de-regulated both during CRP initiation and outgrowth as compared to control. (E) Genes whose expression is de-regulated when CRP progress from initiation and outgrowth stage. (F) Expression pattern of de-regulated genes in different zones of emerged roots. (MZ, meristematic zone; EZ, elongation zone; DZ, differentiation zone).

**Supplementary Fig. S5:** (A) Typical TIC (total ion chromatogram) from GC-MS analysis of amino acids (MSTFA-derivatives) from mock (untreated) and treated (IAA, 12h). Keys to peak identity: 1. L-Alanine, 2. L-Leucine, 3. L-Valine, 4. L-Serine, 5. Isoleucine, 6. Glycine, 7. L-Threonine, 8. L-Aspartic acid, 9. L-5-Oxiproline, 10. L-Phenylalanine 11. L-Glutamic acid, 12. L-Lysine. (B, C) Fold Change (log2 treated vs untreated) of 12 differentially accumulating amino acids from auxin treated samples.

**Supplementary Fig. S6:** Pearson correlation coefficient for rice phosphoproteome data.

**Supplementary Fig. S7:** Representation of percentage of phosphorylation at serine, threonine and tyrosine in differentially up-regulated and down-regulated phosphosites against the total phosphosites.

**Supplementary Fig. S8:** Comparison of de-regulated phosphoprotiens and proteins.

**Supplementary Fig. S9:** GO enrichment analysis of differently regulated phosphosites upon auxin-treatment on the basis of (A) molecular function and (B) cellular localization.

**Supplementary Fig. S10:** Auxin-dependent phosphorylation is involved in cell-differentiation. Heatmap showing dynamic RNA expression pattern of few differentially phosphorylated proteins during CRP initiation and outgrowth. (A, B) Genes specifically de-regulated during CRP initiation (A) or CRP outgrowth (B), with respect to control tissue. (C, D) A few genes are de-regulated during CRP initiation and outgrowth (C), as compared to control tissues or when CRP progress from initiation and outgrowth stage (D). (E) Expression pattern of de-regulated genes in different zones of emerged roots. (MZ, meristematic zone; EZ, elongation zone; DZ, differentiation zone).

**Supplementary Table S1:** GC-MS profile of major amino acids from auxin-treated and untreated rice seedlings. Linear retention index (LRI) was obtained on HP5-MS GC column. Amino acids were identified based on matching with LRI and mass spectrum with standard amino acids/library search. RT: retention time in minutes; LRI cal: linear retention indices calculated from the RT in HP5-MS column using a series of n-alkanes standards (C6–C20); Q-ion: qualifying ion in mass spectra.

**Supplementary Table S2:** A list of primers used in the study.

## Acknowledgements

S.R.Y. acknowledges financial support from Science and Engineering Research Board (SERB), Government of India (grant # ECR/2016/000060). Indian Institute of Technology, Roorkee (IIT Roorkee) is acknowledged to provide infrastructure support to S.R.Y. and fellowships to Z.S. and T.G. Fellowship to H.S. and S.Y. are provided by University Grant Commission (UGC), and Department of Biotechnology (DBT), Government of India, respectively.

## Author Contributions

H.S. and Z.S. have performed experiments of auxin treatment and sample collection for proteomics. H.S. has also performed downstream data analysis, RNA *in situ* hybridization and phenotypic characterization. T.Z. and I.D. have performed (phospho)proteomics related experiments and data analysis. X. X. was involved in analysing the initial test experiments during standardization. B.W. and D.S. performed metabolic profiling and data analysis. T.G. was involved in initial standardization of auxin treatment and spatial expression pattern analysis. S.Y. with H.S. contributed in growing maintaining plants. S.R.Y. conceived the project, designed experiments and analysed data. H.S., T.Z., I.D. and S.R.Y. wrote the manuscript.

## Notes

### Competing Interest Statement

The authors have declared no competing interest.

